# Bioinspired Geometry-Encoded Rheotactic Navigation of Sound-Driven Microrobots

**DOI:** 10.1101/2025.08.31.673395

**Authors:** Chaochao Sun, Adrian Paskert, Yong Deng, Mahmoud Medany, Raphael Wittkowski, Daniel Ahmed

## Abstract

Imitating the shape-encoded tactics of natural microswimmers—organisms that flip, roll, and rheotaxis through viscous fluids—could transform microfluidics, micromanufacturing, and targeted therapy. However, translating those geometric navigation cues into actively driven microrobots is an open, largely unexplored challenge. Here, inspired by the structure of sperm cells, we introduce a sound-propelled head-helix microparticle (“microrobot”) featuring an elliptical head and a spiral tail. This asymmetrical design interacts with the incident acoustic field, generating complex secondary flows that induce a torque, enabling the particle to reorient around its cross-section. The microparticle exhibits a preferred direction of propulsion and orientation when exposed to a traveling sound wave, reorienting if its initial alignment deviates from this preference. Both the preferred direction and orientation can be modulated by adjusting the sound frequency, and they further adapt to background flow fields in the environment. Furthermore, the microparticle exhibits rheotaxis-like motion, exhibiting wall-following motion with frequency-dependent sliding behavior. By moving towards the channel wall, it enters the region with the smallest flow velocities, allowing it to move antiparallel to the fluid. These findings contribute to the engineering of the trajectories of sound-propelled microparticles and to the development of next-generation microrobots for medical and other innovative applications.

## Introduction

Living microswimmers have evolved sophisticated strategies to navigate viscous-dominated environments, employing body geometry, flexible flagella, and non-reciprocal shape deformations to overcome the constraints imposed by the Scallop theorem, as exemplified by bacteria, diatoms, and spermatozoa^1–5^. These locomotion strategies fall into two broad classes. Passive swimmers, such as non-motile diatoms executing Jeffery orbits in shear flow, rely exclusively on hydrodynamics^5–7^. Active swimmers—spirochaetes and sperm—transform biochemical energy into mechanical work, enabling sophisticated trajectories such as upstream migration^2,8,9^. These biological exemplars demonstrate that shape, chirality, and internal actuation can be combined to encode navigation in the absence of external guidance^10^.

Inspired by nature, researchers have created synthetic microparticles that exploit fluid– structure interactions for passive steering^11^. Passive engineered particles exploit fluid-structure interaction to follow various trajectories in complex hydrodynamic fields^12^. Notably, quasi-2D particles with symmetry-broken geometries demonstrate self-steering in confined channels, exhibiting damped oscillatory behavior as they are drawn toward the centerline^13,14^. Structures like “T” or “L” shapes show remarkable predictability in their motion, even under perturbation^15^, and particles with a single mirror plane often trace bell-shaped trajectories^16,17^. Additionally, rough or corrugated channel walls can induce nontrivial trajectories, including controllable helical paths, by modulating hydrodynamic interations^18,19^. Although these designs are predictive, they lack active propulsion, robust autonomy, and controllability—limiting their adaptability and making it difficult to navigate or maneuver effectively in dynamic conditions.

Conversely, active microrobots powered by magnetic fields^20–22^, catalytic reactions^23,24^, light^25–29^, or ultrasound^30–34^ convert external or chemical energy into propulsion and have begun to address biomedical challenges—from clot lysis to biofilm removal and blood‐brain‐barrier translocation. However, most active platforms face a trade-off between propulsion efficiency and trajectory programmability. For example, magnetic helical robots require continuous multi-axis coil steering^21,22^; chemically driven microrobots, such as catalytic Janus particles, rely on self-diffusiophoresis or self-electrophoresis, but their motion is limited by local fuel availability and lacks directional control^23,24^; light-driven microrobots often depend on high-intensity or wavelength-specific illumination, which suffers from limited tissue penetration and line-of-sight constraints^26,27^; and acoustic microrobots typically operate at fixed frequencies, limiting their adaptability to dynamic environments^32,34^. These limitations often necessitate bulky external feedback loops and sophisticated control infrastructures, which are difficult to miniaturize and sustain—particularly within the confined, tortuous spaces of the human body. Thus, a central question remains how to intrinsically encode adaptive navigation into microrobot design, reducing dependence on external intervention while retaining environmental responsiveness.

A promising yet underexplored approach lies in structural encoding—leveraging microrobot geometry to produce adaptive behaviors. Biological systems offer inspiration: Nature provides compelling examples: the intrinsic chirality of sperm tails enables alignment in shear flows, while the asymmetric frustules of diatoms induce passive rotation that enhances nutrient uptake^7,9^. These biological systems demonstrate how structural features alone can generate functional behaviors without external feedback^5,10^. Translating such structure-encoded intelligence to artificial swimmers, however, remains largely unexplored—especially for externally powered, truly active systems.

Among active strategies, acoustically propelled microrobots are particularly promising due to its biocompatibility, safety, deeply penetrating, and already ubiquitous in clinical imaging^32,33^. A diverse range of acoustic microrobots has been developed, achieving translation and rotation through acoustic streaming and radiation forces, often generated by oscillating bubbles or structural appendages^34,35^. In our prior work, we introduced a spiral-shaped acoustic microrobot capable of linear motion^30^; however, the influence of structural geometry on trajectory control and reorientation remained unaddressed. Despite significant progress in acoustic microrobot design, a knowledge gap persists: the critical physics of how geometry couples to acoustic fields to dictate motion trajectories remains poorly understood.

Here, we introduce a sound-propelled microparticle (“microrobot”) with a head-helix design that exhibits geometry-driven behavior in a liquid-filled channel: First, the particle shows a preferred direction of propulsion and a preferred orientation when it is exposed to a traveling sound wave and it reorients if its initial orientation is not in line with its preferred orientation. Second, the preferred direction of propulsion and orientation can be controlled by the sound frequency. Third, they can further change as a reaction to a background flow field in the environment of the microparticle. Fourth, when the microparticle is in a channel, it can be made to move to the channel wall and along it by selecting different sound frequencies over time. This also works if the liquid is flowing through the channel. By moving to the channel wall, the microparticle enters the region with the smallest flow velocities, allowing it to move antiparallel to the fluid. With this rheotactic behavior, the microparticle resembles a property of some microorganisms. These results lay the groundwork for engineering trajectories of sound-propelled microparticles and to the development of next-generation microrobots for medical and other innovative applications.

## Results

### Bioinspired head-helix microrobot

Inspired by the naturally encoded motion trajectories of spermatozoa, we engineered acoustically driven asymmetric microrobots. Each microrobot composed of a spheroidal head and a helical tail (see **Fig. 1**) was fabricated by a high-resolution two-photon lithography 3D printing from a polymeric material IP-S. After 3D printing, the microrobots were developed in propylene glycol monomethyl ether acetate for 15 min to remove the uncured resin. The ellipsoidal head measures 110×90 μm, and the helical tail is 60 μm in diameter and 300 μm in length (see **Methods**). Following fabrication, the microrobots were suspended in a liquid-filled microchannel containing isopropyl alcohol solution with a viscosity of 2.04 mPa·s and a density of 0.785 g/cm^3^. A piezoelectric transducer, bonded adjacent to the acoustic chamber and connected to a function generator, generated acoustic waves to actuate the microrobots (**Supplementary Fig. 1**). A continuous sinusoidal wave was applied at excitation frequencies between 1 to 100 kHz and voltages of 0 to 60 V_PP_. The entire setup was mounted on an inverted microscope, and microrobot motion was recorded using both high-speed and high-sensitivity cameras.

**Fig. 1.**
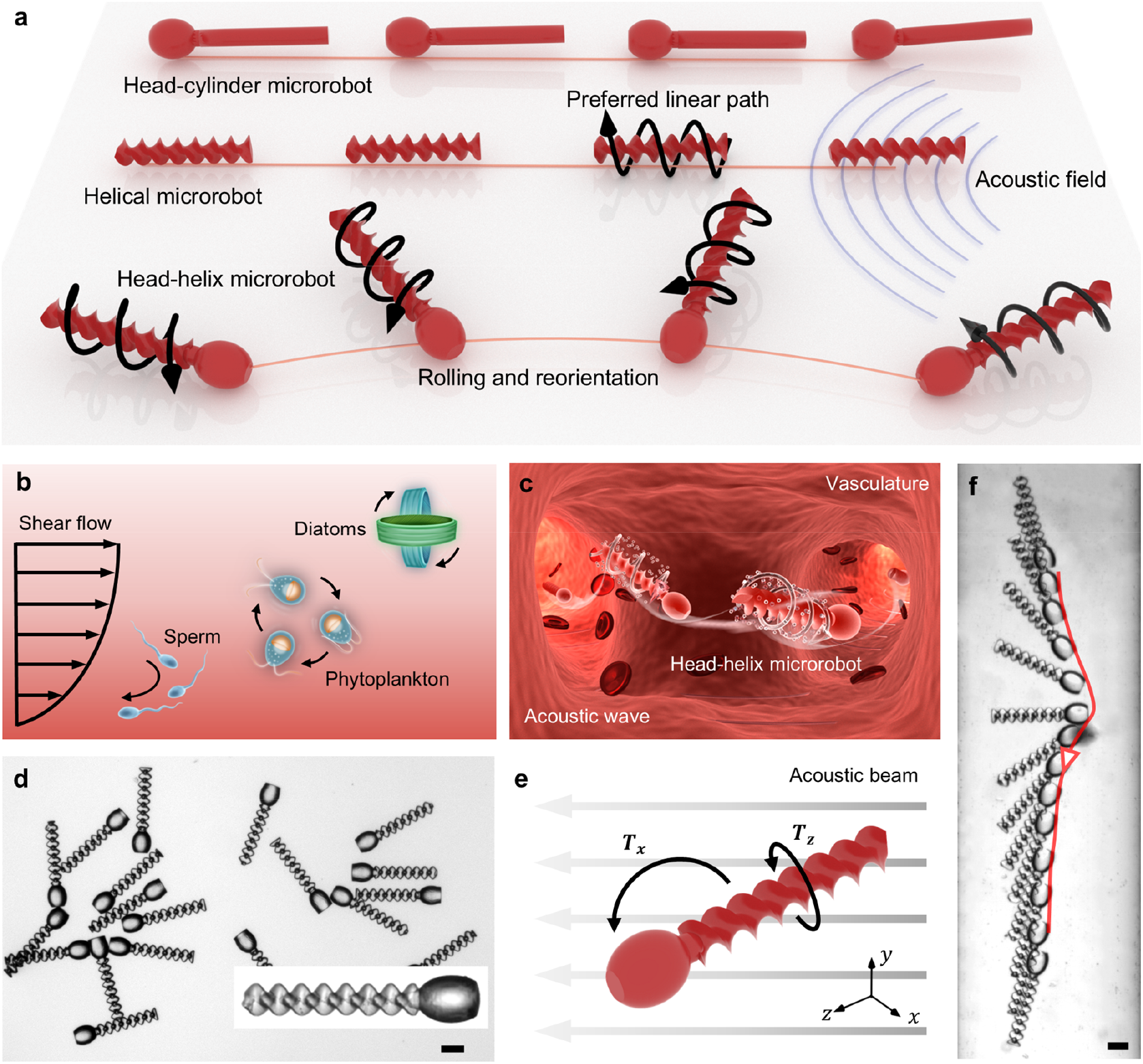
Design and behavior of the head-helix microrobot. **a** Shape-dependent trajectories under an acoustic field, illustrating the distinct motion behaviors of head-cylinder, helical, and head-helix microrobots. The head-cylinder particle exhibits a preferred linear path, the helical particle follows a linear path with axial rotation, and the head-helix particle undergoes both reorientation and axial rotation. **b** Shape-encoded trajectories of various microorganisms under shear flow, illustrating nature’s approach to shape-driven navigation. **c** Concept schematic illustrating the envisioned application of shape-encoded microrobots in complex vasculatures. **d** Fabricated head-helix microparticles. **e** Schematic depiction of the dominant torques acting on the head-helix particle under acoustic excitation. **f** Reorientation trajectory of a head-helix microparticle within a microchannel. Scale bars, 100 μm.

To elucidate how microrobot geometry can intrinsically govern navigation without real-time external feedback, we studied the propulsion and reorientation dynamics of a bioinspired head-helix microparticle subjected to acoustic actuation. Inspired by the helical motility of microorganisms such as spermatozoa, this design enables complex behavior to emerge from structural asymmetry alone. To characterize the behavior of the head-helix microrobot under acoustic actuation, we analyzed its translational motion and reorientation dynamics within a circular microchannel (see **Supplementary Fig. 1b**). Upon excitation, the particle spontaneously begins to roll around its long axis while gradually reorientates its body to align with a preferred orientation relative to the direction of sound propagation. This reorientation reflects a self-regulating trajectory mechanism encoded in the particle’s geometry.

To quantify this reorientation, we define the angle *θ* between the particle’s long axis and the direction of sound propagation. The angular reorientation velocity ω, averaged over the interval from *θ* = 30° to 150°, captures the dominant alignment behavior (**Fig. 2**). At an excitation sound signal of *f*_*1*_ = 12.3 kHz and 60 V_PP_, the asymmetric microstructures reorients from right to left with the tail end leading, reaching an average translational velocity of 259 μm/s and an angular velocity of *ω* = 35.6° (see **Fig. 2a**). Initially, the microrobot is oriented with its head pointing left (0 s), and over the course of 7.4 seconds, it reorients to face right. **Figure. 2b** illustrates the detailed rotation of the microparticle along its long axis, with its rotational direction aligning with the particle’s reorientation direction, which demonstrates the contribution of rotation along the particle axis to the reorientation behavior in the acoustic field.

**Fig. 2.**
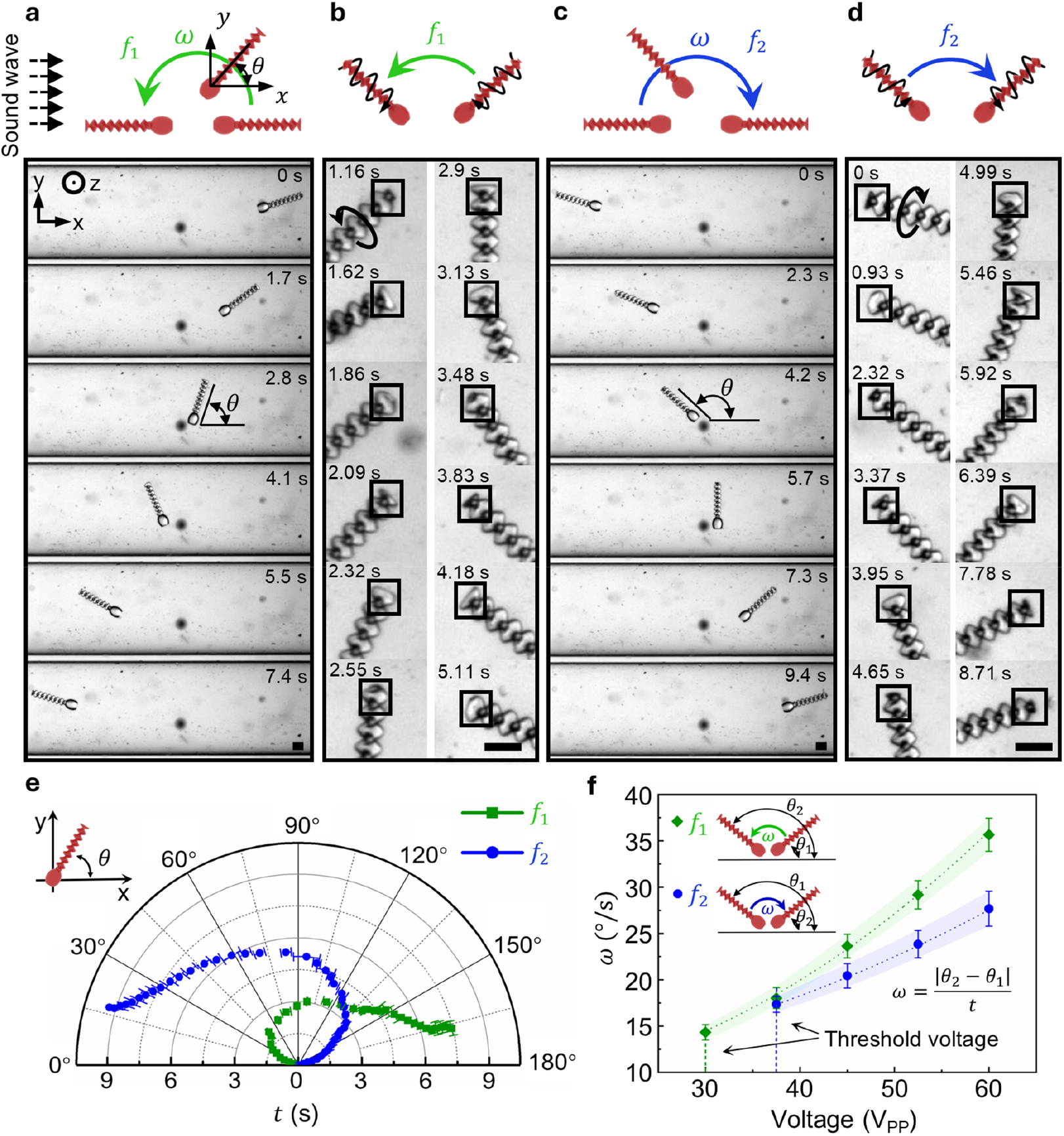
Bidirectional reorientation behavior of the head-helix microrobot in a circular microchannel. **a** Reorientation from right to left in an external acoustic field with *f*_*1*_ = 12.3 kHz and 60 V_PP_. **b** Detailed sequences of the particle rotating around its long axis during the reorientation at *f*_*1*_. **c** Reorientation from left to right in an external acoustic field with *f*_*2*_ = 8.8 kHz and 60 V_PP_. **d** Detailed sequences of the particle rotating around its long axis during the reorientation at *f*_*2*_. **e** The particle’s time-dependent orientation *θ(t)* with time *t* varies with the excitation frequency. **f** The mean angular reorientation velocity as a function of the excitation voltage varies with the frequency. The values were averaged over the angular range from 30° to 150° and fitted with a quadratic function. The threshold voltages show that the particle cannot reorient below 30 V_PP_ at *f*_*1*_ = 12.3 kHz and 37.5 V_PP_ at *f*_*2*_ = 8.8 kHz. Scale bars, 100 μm. The direction of gravity is shown as ☉, i.e., it is perpendicular to the figure plane. Each data point represents the average velocity of at least three measurements. The error bar represents the standard deviation.

Remarkably, this mode of locomotion is fully reversible. At a lower frequency, *f*_*2*_ = 8.8 kHz, the same microrobot reverses direction—both translationally and rotationally—moving from left to right at 218 μm/s and rotating clockwise with *ω* = 27.7°/s (**Fig. 2c**). Besides, **figure. 2d** shows the clockwise rotation of the particle around its long axis at *f*_*2*_ = 8.8 kHz, contributing to the particle’s movement from left to right and its clockwise reorientation. This frequency-dependent bidirectionality reveals that the acoustic field can be used not only for propulsion but also for real-time automatic steer and reverse the particle’s orientation on demand by simply switching the frequency of the acoustic field (see **Supplementary Movie 1**). During this reorientation process, the structure undergoes a reversal in its corkscrew-like motion—i.e., rotation around its longitudinal axis shifts from counterclockwise (CCW; **Fig. 2b**) to clockwise (CW; **Fig. 2d**)— indicating that axial rolling is intrinsically coupled to the swimmer’s reorientation. **Figure 2e** further illustrates the time evolution of the orientation angle *θ* for both frequencies. The particle reorients most rapidly when *θ* approaches 90°, while alignment slows significantly near *θ* ≈ 0° or 180°, suggesting that alignment with the beam axis strongly modulates acoustic torque effectiveness.

To assess the generality of the observed shape-induced reorientation behavior, we investigate whether it depends on channel geometry. Specifically, we examine the microrobot’s motion in an open rectangular microchannel (**Supplementary Fig. 2**) and observe consistent bidirectional behavior—mirroring the dynamics seen in circular microchannels. At an excitation frequency of *f*_*1*_ **=** 13.4 kHz, the particle moves toward the transducer; at *f*_*2*_ = 8.8 kHz, it moves away. In both scenarios, the tail consistently aligns with the direction of net translation, confirming that directional motion is intrinsically guided by the particle’s structural asymmetry rather than by channel confinement.

Although slight frequency shifts are noted between experiments—likely due to minor variations in transducer placement on the glass base plate—the overall behaviors remains robust. Notably, the particle exhibits marginally higher velocities when moving toward the transducer compared to when moving away, a trend also observed in the circular microchannel. This asymmetry may reflect differences in acoustic pressure gradients or streaming profiles near the transducer interface. Together, these results demonstrate that the microrobot’s geometry-encoded navigation strategy operates independently of channel cross-sectional shape. This robustness underscores the versatility of structure-based design for microrobots, enabling predictable and programmable locomotion across diverse fluidic environments without reliance on boundary-guided steering.

To achieve precise and scalable control over microrobot motion, it is essential to understand how propulsion and steering behaviors scale with actuation input. Since the acoustic forces that drive microrobot motion originate from pressure fields generated by the piezoelectric transducer, their strength is expected to depend on the applied voltage. The strength of both acoustic streaming and radiation forces experienced by the microrobot is governed by the intensity of the acoustic field, which is directly controlled by the voltage applied to the transducer. To experimentally validate the relationship, we measure the angular reorientation velocity of the microrobot as a function of excitation voltage. **Figure 2f** demonstrates that the quadratic fit is reasonably well satisfied for the microparticle in both directions. We also observe that the angular velocity at *f*_*1*_ = 12.3 kHz is slightly higher than at *f*_*2*_ = 8.8 kHz, likely due to differences in the resonant amplitude response of the transducer at these frequencies. Importantly, we identify distinct threshold voltages—30 V_PP_ at *f*_*1*_ and 37.5 V_PP_ at *f*_*2*_ and below which no reorientation occurred. This voltage-dependent behavior suggests that the acoustically generated torque, which acts normal to the particle’s long axis, must exceed a critical value to overcome hydrodynamic drag and facilitate reorientation. Since this torque acts primarily normal to the particle’s long axis, the emergence of a voltage threshold highlights the nonlinear nature of acoustic reorientation and propulsion, and underscores the importance of precise control over actuation parameters.

### Shape-encoded reorientation and directional reversal in microrobots

To design microrobots capable of self-guided or adaptive behaviors, it is crucial to understand how structural features contribute to motion control under external fields. In particular, the ability to achieve reorientation—a key behavior for navigation and steering—may depend sensitively on geometric asymmetries. To understand how microrobot geometry governs reorientation under acoustic actuation, we systematically compare three distinct particle designs: a head-helix structure, a pure helix, and a head-cylinder of equal mass to the head-helix. Detailed dimensions of the structures are provided in **Supplementary Fig. 3** and **Supplementary Note 1**. All experiments are performed under identical conditions, using excitation frequencies of *f*_*1*_ **=** 12.3 kHz, *f*_*2*_ **=** 8.8 kHz, and a driving voltage of 60 V_PP_, with acoustic waves propagating from left to right. Our goal is to determine which features are necessary and sufficient to enable geometry-induced reorientation under acoustic actuation.

As shown in **Fig. 3a**, the head-helix microrobot exhibits frequency-dependent bidirectional reorientation in response to acoustic stimulation. This behavior is characterized by the microrobot aligning its helical tail along the propulsion direction following a torque-induced reorientation. In contrast, the helix-only particle—lacking a head—undergoes frequency-dependent reversal of translational motion (at *f*_*1*_ **=** 12.3 kHz, and *f*_*2*_ **=** 8.8 kHz, **Fig. 3b**), but does not exhibit any body reorientation. While the helical geometry enables corkscrew motion and propulsion, the absence of a front-end asymmetry removes the torque necessary for out-of-plane rotational reorientation.

**Fig. 3.**
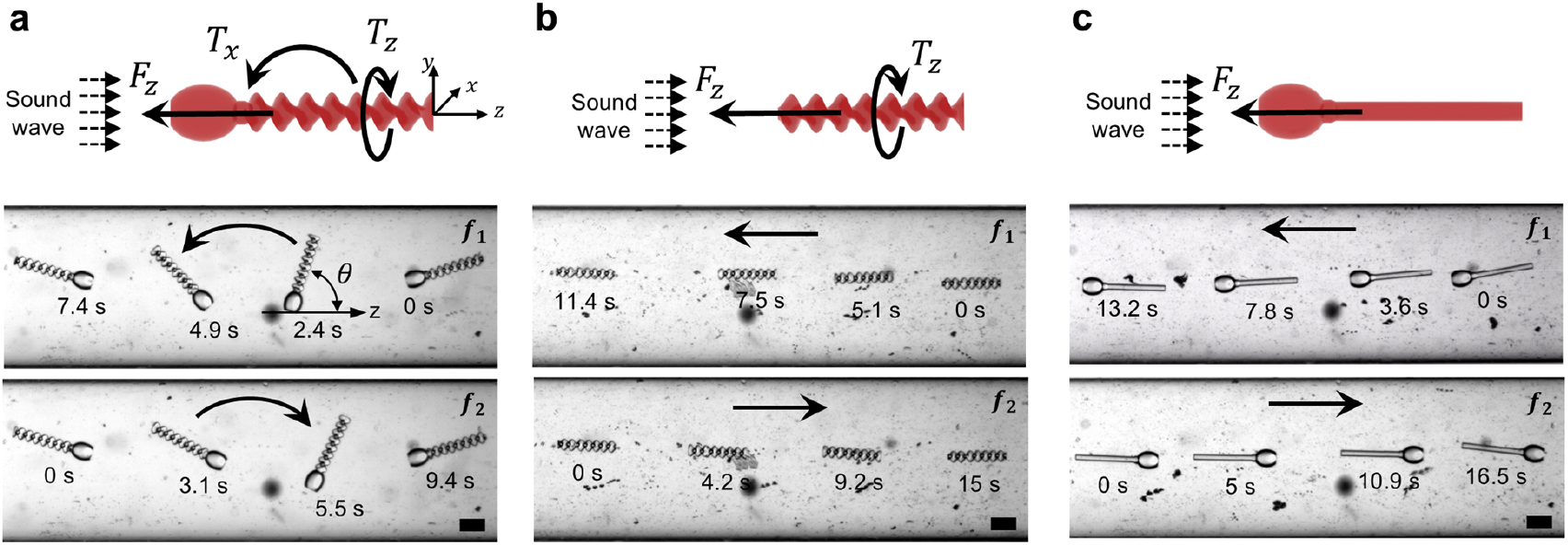
Shape-dependent bidirectional motion behavior of three types of microrobots. **a** Bidirectional reorientation of the head-helix particle. **b** Bidirectional translational corkscrew motion of the helical particle without reorientation. **c** Bidirectional translational motion of the head-cylinder particle without reorientation. The sound travels from left to right in all cases (*f*_*1*_ = 12.3 kHz, *f*_*2*_ = 8.8 kHz, 60 V_PP_). Scale bars, 100 μm.

We then examine head-cylinder particles, which possess asymmetry but lack a helical tail. These microparticles also translate in both directions depending on the excitation frequency, similar to the other designs (**Fig. 3c**). However, no reorientation is observed, even when the initial orientation is varied. This suggests that while the head may contribute to directional propulsion, it does not generate sufficient torque to overcome hydrodynamic drag and induce reorientation in the absence of a chiral tail. Together, these results highlight that both structural asymmetry (from the head) and chirality (from the helix) are necessary to generate the complex torque profiles required for reorientation.

Additionally, we observe that regardless of the initial orientation of the head-helix particle, its final orientation consistently features the helix leading the particle in the propulsion direction (see **Supplementary Fig. 4a-b**). This alignment indicates that the helical tail generates a stronger propulsion force than the head, driving the reorientation process. This conclusion is further supported by showing that the helical particle achieves a higher translational velocity than the head-cylinder particle, even though the latter has a larger overall volume (**Fig. 3b** and **3c**). The enhanced propulsion of the helical particle reinforces the dominant role of the tail in sound-driven motion. However, the head-cylinder particle moves independently of its initial orientation and does not exhibit any reorientation behavior, as shown in **Supplementary Fig. 4c-d**. These findings highlight that adjusting the initial orientation of the head-helix particle can also enable tailored path-planning strategies, providing versatile control in navigating complex environments.

Although we have previously established that the head–tail asymmetry and helical-tail structure enable reorientation, there remains a lack of direct evidence to fully explain why the head-cylinder particle does not exhibit reorientation under similar conditions. Despite having a tail of similar length and volume to the helical tail, the head-cylinder particle does not reorient, regardless of whether the initial orientation is *θ < 0* or *θ > π*. We therefore hypothesize that the helical geometry of the tail plays a key role in enabling reorientation by inducing axial rotation with a fixed handedness, which breaks symmetry and drives rolling during the reorientation process.

To further elucidate the role of acoustic streaming in the reorientation behavior of the head-helix microrobot, we investigated the acoustic streaming distribution and computed the axial acoustically induced torque through acoustofluidic computer simulations. Since the streaming effects that manifest around the particle are inherently caused by the non-linearity of the underlying fluid dynamics, direct numerical simulations of the compressible Navier-Stokes equations are needed. The momentum-stress tensor *∑*_*ji*_ can be expressed as

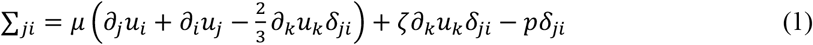

Here, *t* denotes time, *∂*_*i*_ is the partial derivative with respect to the *i*th spatial coordinate, *u*_*i*_, *ρ*, and *p* are the velocity, mass density, and pressure fields. The viscosity of the fluid is modeled through the constant shear- and bulk viscosities *μ* and *ξ* and *δ* is the Kronecker-delta symbol.

Because the stress tensor contains the force density exerted on the particle by the fluid, integration of this quantity over the surface of the particle yields the total force exerted on the particle. By considering the position *r*_*i*_ of each force element relative to the microparticle’s center of mass, the torque density can be written as ∈_*ij*_ *r*_*i*_*n*_*i*_*∑*_*j*_ where ∈ is the Levi-Civita symbol and *n*_*i*_ denotes the unit outward normal vector to the surface. The total torque *T*_*i*_ on the microrobot can be expressed as an integral over the microrobot’s surface as

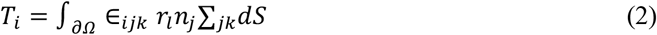

Microrobot dynamics were simulated using the same system and parameter values as in the experiments by numerically solving the governing equations for sound propagation in the fluid and its interaction with the particle (see **Methods** and **Supplementary Note 4**). The acoustofluidic simulations revealed significant acoustic streaming around the head-helix microrobot, which is consistent with the magnitude observed in the experiments (**Supplementary Movie 2**). The streaming flow is concentrated near the fins of the helical tail, where it generates a rotational component that induces torque along the particle’s long axis. A schematic analysis illustrates how the axial torque contributes to the reorientation process, with the direction of this torque depending on both the excitation frequency and the initial orientation (θ = 0 or π), as shown in **Fig. 4a**. The simulated torque values for four representative cases are summarized in **Fig. 4b**. For frequency *f*_*1*_, torques of –88 μN·μm at θ = 0 and 15 μN·μm at θ = π drive counterclockwise (CCW) reorientation. In contrast, For *f*_*2*_, the torques are reversed (–63 μN·μm at θ = π and 35 μN·μm at θ = 0), resulting in clockwise (CW) rotation. These simulation-based predictions are strongly supported by experimental observations of the surrounding flow field using particle image velocimetry (PIV). At *f*_*1*_, the microrobot exhibits CCW acoustic streaming patterns, while at *f*_*2*_, CW streaming dominates (**Fig. 4c, d**; **Supplementary Movie 3**). Moreover, the observed reorientation behaviors of the microrobot (**Fig. 2b, d**) align closely with the predicted torque directions in all four scenarios, confirming the robustness of the acoustofluidic model. These findings collectively confirm that the helical tail structure, by inducing a strong rotational flow field and torque along the particle’s long axis, plays a pivotal role in driving the reorientation behavior of the head-helix microrobot.

**Fig. 4.**
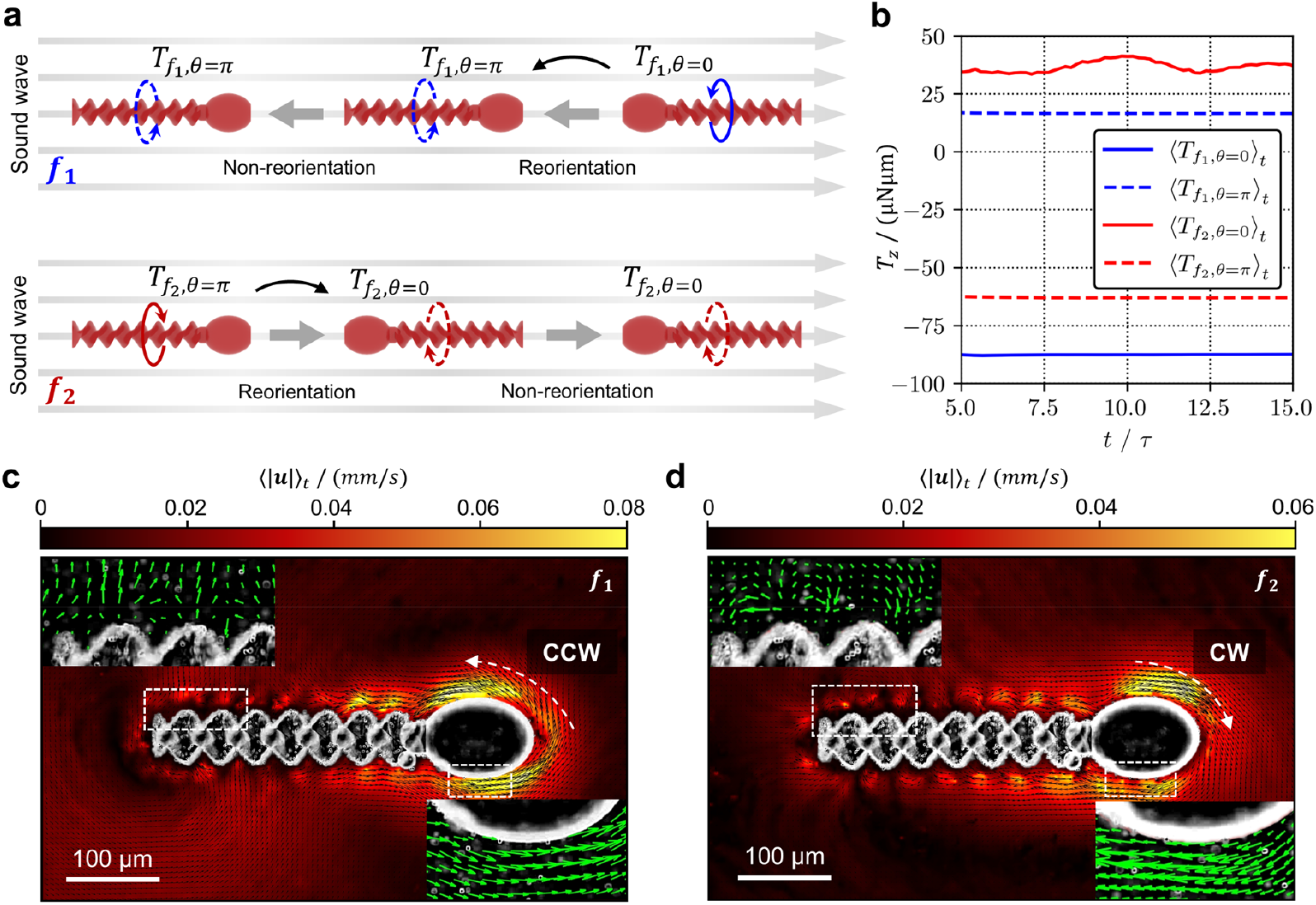
Torques and acoustic streaming patterns at different excitation frequencies. **a** Schematic illustrating the motion behaviors of the head–helix microrobot and the associated torques under different acoustic frequencies and orientations. **b** The torque values acting on the microrobot in four different scenarios, obtained from acoustofluidic simulations using the fluid-dynamics solver AcoDyn. **c-d** Experimentally measured flow fields visualized by particle image velocimetry (PIV) at two excitation frequencies, *f*_*1*_ and *f*_*2*_, respectively. The resulting acoustic streaming patterns are asymmetric, with counterclockwise (CCW) vortices at *f*_*1*_ (**c**) and clockwise (CW) vortices at *f*_*2*_ (**d**). Prominent flow circulations are observed near the microrobot’s head and along the helical tail. Scale bar, 100 μm.

### Reorientation behavior in the presence of sound and external flow

Understanding the influence of external flow on particle motion within an acoustic field is critical for practical applications where flow and acoustic forces coexist. Accordingly, we examine the behavior of head-helix particles under four distinct conditions—low and high flow rates in the absence of sound, and no flow and low flow rates in the presence of sound—to elucidate the combined effects of hydrodynamic and acoustic forces on particle dynamics (see **Fig. 5a-d**). First, the particle is placed into the channel in a countercurrent orientation. Upon turning on the pump, the particle rapidly moves at high speed and shows reorientation motion only driven by a high flow field (*u* = 1660 μm/s) (see **Fig. 5a**). This behavior is attributed to the interaction between the helical structure and the gradient flow within the channel, which causes the helical structure to drift perpendicular to the shear flow at low Reynolds numbers^36,37^. The whole process occurred within 1.5 seconds. Next, when the flow rate was set at a low flow rate (*u* = 330 μm/s), the particle moves along the channel without reorientation (see **Fig. 5b** and **Supplementary Movie 4**). For the particle, there is a threshold flow rate below which it cannot reorient, resembling the behavior of sperm, which also encounter difficulties in reorienting at low flow rates^38^. **Figure 5c** illustrates that our design exhibits reorientation behavior again if, besides the flow field, sound is present, even when the flow rate is low (*u* = 330 μm/s, *f*_*1*_ = 12.3 kHz, 60 V_PP_) (also see **Supplementary Movie 4**). Notably, the particle achieves reorientation behavior in just 2.6 seconds, which is comparable to the case where it is exposed only to a high flow rate and faster than for the case where it is exposed only to a sound field (taking 4.9 seconds to realize reorientation), as shown in **Fig. 5d** (*f*_*1*_ = 12.3 kHz, 60 V_PP_). Thus, these cases comprehensively demonstrate that our designed head-helix particle can reorient in response to both external fields, whether it is a flow field or a sound field.

**Fig. 5.**
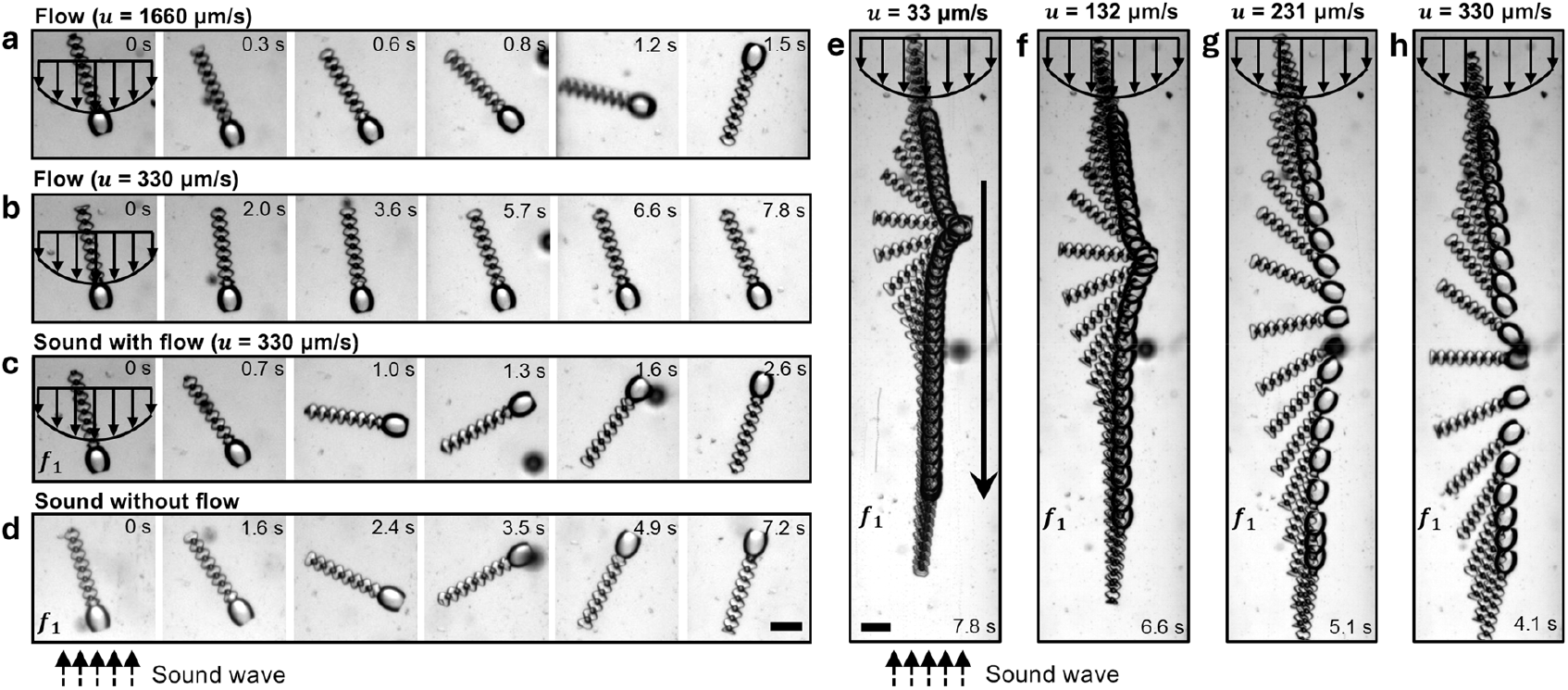
Reorientation behavior of the head–helix microparticle under sound and external flow fields. **a** Reorientation at a high flow rate of *u* = 1660 μm/s. **b** Absence of reorientation at a low flow rate of *u* = 330 μm/s. **c** Reorientation driven by combined sound and flow fields at *u* = 330 μm/s, *f*_*1*_ = 12.3 kHz, and 60 V_PP_. **d** Reorientation in a sound field without external flow at *f*_*1*_ = 12.3 kHz and 60 V_PP_. **e-f** Reorientation at different flow rates ranging from 33 μm/s to 330 μm/s under *f*_*1*_ = 12.3 kHz and 60 V_PP_. Scale bars, 100 μm.

We then characterize the effect of the flow rate on the particles’ trajectory in the case of combined sound and flow fields. **Figure 5e-h** shows stacked images of a particle at different time points during its reorientation motion in the combined case, with an interval of 0.233 seconds between two adjacent images. As the flow rate increases, the driving force acting on the microparticle intensifies, providing it with greater kinetic energy and leading to high propulsion velocity. With the increase of the flow rate, the maximum angle difference between adjacent images increases, indicating a higher maximum reorientation angular velocity *ω*. It can be concluded that the increased flow rate leads to a higher torque *T*_*x*_, which accelerates the overall reorientation process. Thus, we demonstrate that precise trajectory control of head-helix particles can be achieved through regulation of both acoustic and flow fields. This dual-field control mechanism holds significant application value in the field of targeted transport for microrobot.

### Rheotactic behavior in a sound field

Propulsion and upstream motion represent fundamental behaviors observed in natural microswimmers, as exemplified by spermatozoa navigating in shear flows (**Fig. 6a**). These biological systems achieve rheotaxis—the ability to orient and swim against fluid flow^9,39^. A similar behavior can be observed with our head-helix particles. **Figure 6b** demonstrates frequency-dependent rheotactic behavior of our particles in quiescent fluid under two distinct acoustic excitation frequencies. At first, the particle is in the center of the channel and exhibits reorientation in response to the excitation frequency *f*_*1*_ = 12.3 kHz. Subsequently, the particle starts to move toward the wall at 2.8 seconds and exhibits propulsion motility along the wall as the frequency is switched to a value *f*_*3*_ = 18.6 kHz (see **Supplementary Movie 5**). In the process of propulsion, the particle advances unidirectionally along the channel at a constant speed while also rotating about its axis. This movement is balanced by gravity, acoustic radiation force, hydrodynamic interaction of the rotating particle with the surface, and drag force as it reaches the channel wall (see **Supplementary Fig. 5** and **Supplementary Note 2**). To investigate the effect of the excitation voltage on its propulsion, the propulsion velocity is measured versus excitation voltage, as shown in **Fig. 6d**. The results show that the propulsion velocity *v*_*t*_ also scales quadratically with the applied voltage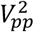, which is governed by acoustic radiation forces.

**Fig. 6.**
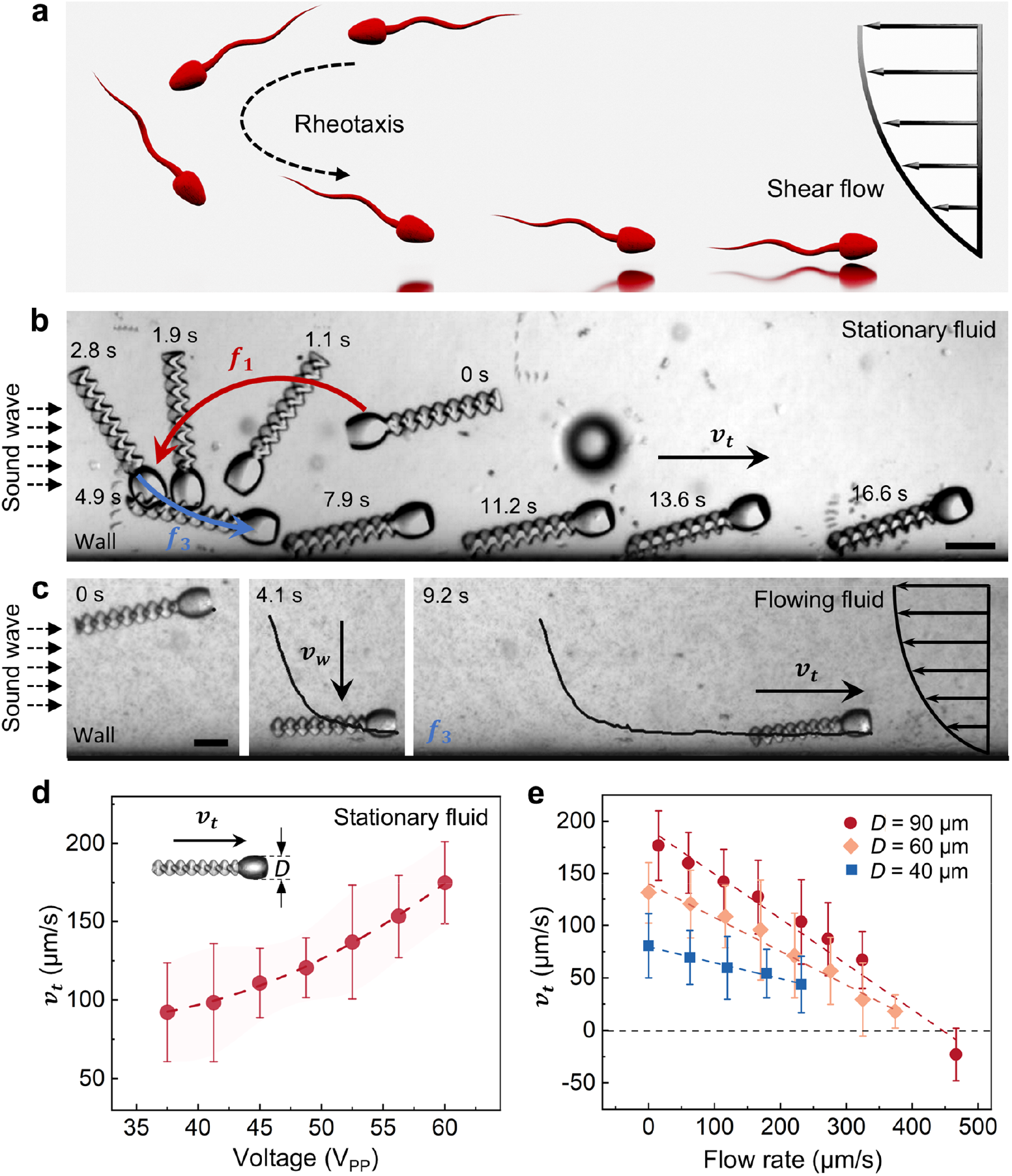
Rheotactic behavior in a sound field. **a** Rheotaxis of a spermatozoon in shear flow. **b** Rheotactic behavior of the head-helix particle in a sound field at *f*_*1*_ = 12.3 *kHz, f*_*3*_ = 18.6 kHz and 60 V_PP_. *v*_*t*_ denotes the propulsion velocity of the microrobot along the wall at *f*_*3*_. **c** Image sequences demonstrating the upstream motion of the particle (*D* = 90 μm) along the channel wall in an sound field. The trajectory is shown together with the oppositely oriented flow field. Scale bar, 100 μm. **D** Propulsion velocity *v*_*t*_ of the particle (*D* = 90 μm) as a function of acoustic driving voltage in stationary fluid. **e** Propulsion velocity *v*_*t*_ as a function of channel flow rate. All measurements in **c-e** were performed at *f*_*3*_ = 18.6 kHz and 60 V_PP_.

Under physiological conditions in living systems, such as humans, the preeminent obstacle is the blood flow velocity. To execute all kinds of tasks in vivo, particles should be able to resist the flow and navigate the flowing fluid. Thus, the upstream ability of the particle has been tested in a straight channel. **Figure 6c** depicts the propulsion process of microparticles within a flowing fluid when excited at frequency *f*_*3*_. A particle rotating around its axis exhibits left-to-right propulsion when it reaches the wall, which is traced with a black trajectory. The symbols *v*_*w*_ and *v*_*t*_ represent the particle’s velocity from the channel’s center to the wall and propulsion velocity along the wall, respectively. Given the gradient flow in the channel, particle size is expected to strongly influence behavior. To examine this effect, we analyze three particles of the same shape but different head diameters (*D* = 40, 60, and 90 μm; see **Supplementary Fig. 6**). While *v*_*w*_ remains almost unchanged with increasing flow rate, larger particles attain higher velocities owing to the stronger radiation force, which scales with particle volume (see **Supplementary Fig. 7** and **Supplementary Note 3**). **Figure 6e** shows the upstream velocity *v*_*t*_ of three particles against the flow rate. We consider the average upstream velocity for at least five measurements for each flow rate. The results indicate that the upstream velocity *v*_*t*_ decreases as the flow rate increases. Remarkably, the particle (*D* = 90 μm) is able to move upstream against a flow rate as high as 324 μm/s, despite exhibiting a maximum propulsion velocity of only 177 μm/s in the stationary fluid under identical acoustic excitation parameters, beyond which it is swept away by the flow (see **Supplementary Movie 6**). The propulsion velocity *v*_*t*_ of the particles decreases linearly for increasing flow rate^31^, which is satisfied reasonably by the microparticle rolling along the wall. The slight deviation of the linear fit is expected given by the various distribution of the microparticle’s diameter during rheotaxis.

## Discussion

Our results demonstrate that this design enables a controlled bidirectional reorientation behavior that can be tuned by the sound frequency. The translational and rotational motion enables the particle’s ability to mimic rheotactic motion, akin to natural microswimmers like sperm cells, demonstrates that carefully engineering the shapes of motile microparticles has the potential to design microparticles whose trajectories can reach a similar complexity as the trajectories of microorganisms.

The observed rheotactic behavior of the microparticles aligns with the natural movement of sperm cells, suggesting that bio-inspired structural designs can effectively replicate biological functionalities. The ability to control orientation and directionality through sound frequency modulation represents a novel advancement beyond natural systems. This reorientation behavior, which is not observed in previous acoustically actuated microparticles, underscores the significance of shape rather than material design and complex external control systems in achieving complex motion profiles. The unexpected finding that the particle (*D* = 60 μm) can resist fluid flow rates up to 374 μm/s, 2.6 times their propulsion velocity of 141 μm/s in stationary fluid, demonstrating a level of flow-field interaction previously unattainable in synthetic systems. This capability also opens new possibilities for biomedical applications in complex fluid environments, such as microsurgical tools that adjust their trajectory in real-time to navigate intricate ductal networks (e.g., biliary or bronchial systems) and enhanced diagnostic sampling like biomarkers by programming particles’ reorientation.

While prior studies have explored magnetic, optical, and chemical actuation methods for microrobots, these approaches often face limitations such as low penetration depth, complex equipment, and biocompatibility issues. In contrast, our use of acoustic fields offers a non-invasive and portable alternative. Additionally, translational velocities increase quadratically with input voltage, allowing significant enhancement of particle propulsion by increasing voltage. Our findings extend this knowledge by demonstrating that acoustic fields can enable precise control over reorientation and rheotactic behavior, which has not been achieved with other actuation methods.

Harnessing sound fields to control microparticle trajectories introduces transformative opportunities in biomedical applications such as targeted drug delivery, minimally invasive surgeries, and enhanced assisted reproductive technologies (ART). These particles navigate upstream against fluid flows, enhancing both the targeted delivery of genetic materials like sperm DNA and direct egg interaction—paving the way for more effective and controlled fertilization. The particles’ design allows for customization in size and dual functionality, where drugs can be loaded both at the head and tail. This innovation enables the development of sophisticated “smart” drug delivery systems. Equipped with pH-sensitive or temperature-sensitive polymers, these systems release drugs in a controlled manner, responsive to environmental conditions. This targeted approach is crucial for treating complex diseases like cancer, where precision and adaptability in treatment methods substantially improve patient outcomes. Moreover, the integration of acoustofluidic systems with lab-on-a-chip technologies significantly advances microfluidics and micromanufacturing. These advancements are crucial for creating efficient platforms for diagnostics, chemical synthesis, and environmental monitoring, and they play a pivotal role in neurological applications such as deep brain stimulation and targeted drug delivery.

While this study emphasizes the importance of particle design in achieving programmable trajectories, a critical focus area moving forward is the topology optimization to enhance reorientation and fluid navigation capabilities. Future research should prioritize exploring the interactions between sound-propelled microparticles of other various shapes and dynamic flow fields. Overcoming these challenges will involve leveraging computational simulations, machine learning, and innovative designs with intricate shapes and specific trajectories to drive significant advancements. Additionally, developing multifunctional particles responsive to diverse stimuli like sound and magnetic fields, along with integrating materials that respond to environmental cues for precise drug release control, will enhance “smart” drug delivery systems, leading to substantial progress in medical treatments.

In conclusion, our findings demonstrate the potential of bio-inspired, acoustically controlled microparticles for precise trajectory manipulation in complex environments. By addressing current limitations and exploring future directions, this work paves the way for the development of next-generation microrobots with transformative applications in medicine, manufacturing, and beyond. The unique reorientation and rheotactic capabilities of our design represent a significant step forward in the field of microrobotics, offering new opportunities for innovation and practical implementation.

## Methods

### Fabrication of microparticles

The head-helix microparticles were designed using 3D Max software. DeScribe was used to generate the job program for the 3D printing, and default parameters were adopted for the model preparation. A commercial two-photon polymerization system (Photonic Professional GT, Nanoscribe GmbH, Karlsruhe, Germany) was adopted to print the microparticles with a 25× objective. The microparticles were printed on the ITO glass substrates using high-resolution photoresist IP-S (Nanoscribe GmbH, Karlsruhe, Germany). The photoresist was cured programmatically until the job was done. After 3D printing, the microparticles were developed in propylene glycol monomethyl ether acetate (PGMEA, Sigma-Aldrich Inc., St. Louis, MO, USA) for 15 min to remove the uncured resin.

### Acoustic excitation setup

The experiments were carried out in a circular microchannel (800 μm in diameter) or an open rectangular channel (10 mm × 10 mm) in PDMS (Dow Corning). Different straight microchannels (800 μm in diameter), including 45° and 90° tilt angles, were fabricated. A glass slide (50 mm × 25 mm × 1 mm) coupled with such a PDMS microchannel and a transducer (Steiner & Martins, Inc., USA) was mounted on an inverted microscope (Carl Zeiss, Germany) with various objectives (2.5x, 5x, 10x, 20x) for imaging, as shown in **Supplementary Fig. 1**. The excitation system mainly consists of a function generator (Tektronix, Inc., USA) and an amplifier (Digitum-Elektronik, Germany). A continuous square wave was applied to the piezo-transducer. A standard camera (Photometrics, USA) and a high-speed camera (Kron Technologies Inc., Canada) were used to record the time-lapse images for further analysis. Isopropyl alcohol solution was injected into either an open rectangular channel or a circular microchannel as the working fluid, after which microparticles were introduced via a microfiber.

### Upstream motion

When demonstrating a microparticle’s upstream motion driven by an external sound field, the flow rate was regulated by a programmable syringe pump connected to the microchannel. The average flow rate was determined by injecting a constant volume of fluid into the microchannel and measuring the time required for the fluid to travel a specified distance along the channel. To minimize the impact of varying friction resulting from liquid accumulation on the channel walls, the experiment was performed with a controlled volume (instead of a fixed pressure). A solution consisting of 10:1 by volume IPA and 2.0 μm tracer particles (Polysciences, USA) was introduced inside the silicon tube of the syringe pump. The upstream motion was recorded by a camera and analyzed using software ImageJ.

### Simulation method

The simulation results were obtained using the in-house developed finite volume method simulation framework AcoDyn (version 3.1). For the simulation setup, a cylindrical fluid domain with a radius of 400 μm was created around the microparticles (for detailed dimensions, see **Supplementary Fig. 3**). The properties of the compressible fluid surrounding the particle were modeled after water at room temperature, with a shear and bulk viscosity of *μ*_*s*_ = 1.002 mPa s and *μ*_*b*_ = 2.87 mPa s, respectively. The normal density at atmospheric pressure was set to *ρ*_*0*_ = 998.2 kg/m^3^ whereas the speed of sound was set to a constant *c* = 1484 m/s. A traveling wave was generated using a time-periodic boundary condition on the cylinder face behind the microparticle and damped using a time-periodic forcing zone on the opposite cylinder face to prevent unwanted reflections. The pressure amplitude of the generated wave was set to *Δ* = 60 kPa, with a corresponding velocity amplitude of approximately 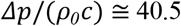 mm/s. For the mantle of the cylindrical fluid domain, as well as on the particle surface, we prescribed no-slip boundary conditions. The simulations were run to near convergence at 30-50 wave periods. To ensure accurate results, the fluid domain was discretized to an unstructured tetrahedral grid with an inhomogeneous resolution of 3.5 to 4.5 million finite volume cells, depending on the specific simulation run, where the largest grid refinement was always located near the microrobot’s surface. Satisfying the Courant-Friedrichs-Lewy criterion based on the speed of sound in the medium^40,41^, the time-step size was limited to 0.98 ns. To limit numerical diffusion due to floating point inaccuracies, the simulation was conducted on non-dimensionalized quantities, with characteristic values for the velocity *v*_*c*_ = 40.5 mm/s, pressure 𝒫_*c*_ = 60 kPa, density *ρ*_*c*_ *= ρ*_*0*_ = 998.2 kg/m^3^, and length *(σ* = 1 μm.

## Supporting information

manuscript

## Data availability

The authors declare that data supporting the findings of this study are available within the paper and its Supplementary Information. The source data can be provided by the corresponding author upon reasonable request.

## Author contributions

D.A. conceived the project and supervised C.S. and Y.D.. R.W. supervised A.P.. C.S., Y.D., and D.A. contributed to the experimental design. S.C. performed the experiments. A.P. performed the computer simulations. A.P. and R.W. contributed to the theoretical understanding. S.C., A.P., Y.D., M.M., R.W., and D.A. contributed to data analysis, scientific presentation, discussion, and manuscript writing.

## Acknowledgements

This project has received funding from the European Research Council (ERC) under the European Union’s Horizon 2020 research and innovation program grant agreement no. 853309(SONOBOTS) and ETH Research grant ETH-08 20-1. R.W. is funded by the Deutsche Forschungsgemeinschaft (DFG, German Research Foundation) — 535275785. The simulations for this work were performed on the GPU nodes of the computer cluster PALMA II of the University of Münster. C. S. acknowledges the financial support from the China Scholarship Council (202206020125). C. S. extends sincere gratitude to Prof. Songmei Yuan of Beihang University and Prof. Jian Lu of City University of Hong Kong for their unwavering support throughout this work.

