## Supplementary material for "Bioinspired Geometry-Encoded Rheotactic Navigation of Sound-Driven Microrobots": manuscript

**This PDF File Includes:**

Supplementary Notes 1 to 4

Supplementary Figs. 1 to 7

References

### Supplementary Note 1. The design of the detailed dimensions of microparticles

To investigate the primary contribution resulting in the reorientation of the microparticles, three types of topological structures were designed: head-helix, helical, and head-cylinder particles. To ensure the head-cylinder particle possesses the same mass asymmetry as the head-helix particle while minimizing the influence of mass differences, the tail volume of the head-helix particle was calculated using SolidWorks software, yielding a value of $2.58 \times 10^{-4}$ $mm^3$. Based on this calculation, the tail diameter of the head-cylinder particle was determined to be 32 μm (see **Supplementary Fig. 3**).

### Supplementary Note 2. The hydrodynamic model for the upstream particle in a circular channel

In the context of the movement of a head-helix microparticle through a circular channel under the influence of an external acoustic field, a combination of forces is exerted upon the particle, including gravity $F_G$ and acoustic radiation forces $F_A$[1,2]. The spiral structure of the microparticle's tail induces acoustic streaming, thereby generating a net torque $T_Z$ that drives the particle to rotate along its longitudinal axis. As the head rolls near a surface, it encounters a net resistive force $F_R$ due to fluid interactions, creating additional torque that contributes to the curved trajectory[3]. This interaction gives rise to a lateral force, which is further influenced by hydrodynamic effects. Similar to the motion of natural sperm cells, the head-helix particle experiences a drag force $F_D$ that acts in opposition to its propulsion, arising from fluid resistance during its swimming-like motion[4]. The propulsion of the particle is governed by a balance of forces, including gravitational forces, acoustic radiation forces, lubrication forces, and drag forces, which stabilize its movement as it approaches the channel wall (see **Supplementary Fig. 5**). Consequently, the particle achieves unidirectional propulsion along the channel at a steady velocity while maintaining continuous rotation around its axis.

### Supplementary Note 3. Shift velocity of the particles

The effects of flow rate and particle size on the shift velocity ($v_w$) of the particles moving from the channel center to the wall were investigated under conditions of $f_3 = 18.6$ kHz at 60 $V_{PP}$. The average shift velocities were measured and analyzed using the ImageJ tracking plug-in. The average flow rate was determined by injecting fluid into an empty microchannel and calculating it based on the fluid length over a specific time interval. As particle size increases, the shift velocity ($v_w$) also increases, as the radiation force, which is proportional to the particle's volume, dominates this process[5,6]. Results for all particle sizes indicate that the shift velocity increases slightly with higher flow rates. The gradient flow distribution enhances the movement of asymmetric particles toward the channel wall, a behavior resembling rheotaxis[4]. As shown in **Supplementary Fig. 7**, the trajectory's curvature is large at minimal flow rates, while it approaches a right-angle trajectory at higher flow rates.

### Supplementary Note 4. Acoustofluidic simulations

In order to accurately assess the streaming and pressure field around the microparticle, acoustofluidic simulations were conducted. Since the streaming effects that manifest around the particle are inherently caused by the non-linearity of the underlying fluid dynamics, direct numerical simulations of the compressible Navier-Stokes equations are needed. As the volume of the simulated fluid must be kept small enough to ensure that the computational effort remains feasibly with the available computational resources, the experimental setup was approximated with the particle placed in the middle of a cylindrical channel. The acoustic wave is then induced into the fluid domain through one cap of the fluid domain such that it travels in parallel to the axis of the cylindrical domain.

The structure-fluid interaction between the particle and the surrounding liquid was modeled using an impenetrable no-slip boundary condition. Modeling a particle at rest, this allows us to study particle-fluid interactions without having to consider the dynamics of the structure itself. This is justified as the material of the microparticle used in the experiments has a sufficient rigidity. Since the velocity of the particle is orders of magnitude lower than the fluid velocity of the surrounding oscillating field, simulating a static particle is a good approximation. A similar boundary condition was employed for the mantle surfaces of the channel around the microparticle, as this models the setup in the experiments and avoids numerical instabilities.

To generate the acoustic wave, Dirichlet boundary conditions were used on the inlet side of the cylinder where the pressure- and velocity values were modulated following

$$p = p_0 sin(tf) \tag{1}$$

$$u = u_0 sin(tf) \tag{2}$$

were $p_0 = 60$ kHz is a pressure amplitude and $u_0 = \frac{p_0}{\rho_0 c_f}$ $i$s the velocity amplitude with a normal density of water at room temperature of $\rho_0 \approx 998.2\ kgm^{-3}$ and a speed of sound of $c_f = 1484\ ms^{-1}$. For the density at this inlet, a zero-gradient von Neuman boundary condition was utilized. To avoid unwanted reflections at the other end of the channel and back at the microparticle, a forcing zone was implemented that efficiently absorbs the wave with minimal reflections.

In addition to the Navier-Stokes equations, the continuity equation, as well as a thermodynamic state equation were used as the governing equations to model the fluid flow. The numerical implementation was realized entirely within the self-developed finite volume solver package AcoDyn. From the numerically calculated time-averaged density, velocity, and pressure fields, the structure of the streaming field can be studied, and the net propulsion force and torque exerted on the particle surface by the ultrasound wave can be calculated. To obtain the long-time solution of the fluid dynamics around the particle, each simulation is run until the time-averaged fields are sufficiently converged to a steady state.

To derive the propulsion force from the numerical field values, the Navier-Stokes equations are written as

$$\frac{\partial}{\partial t}\rho u_i + \partial_j \rho u_j u_i = \partial_j \Sigma_{ji} \tag{3}$$

where the momentum-stress tensor $\Sigma_{ji}$ can be expressed as

$$\Sigma_{ji} = \mu\left(\partial_j u_i + \partial_i u_j - \frac{2}{3}\partial_k u_k \delta_{ji}\right) + \zeta \partial_k u_k \delta_{ji} - p\delta_{ji} \tag{4}$$

In this notation, $t$ denotes time, $\partial_i$ is the partial derivative with respect to the $i$th spatial coordinate, $u_i$, $\rho$, and $p$ are the velocity, mass density, and pressure fields. The viscosity of the fluid is modeled through the constant shear- and bulk viscosities $\mu$ and $\zeta$ and $\delta$ is the Kronecker-delta symbol.

Because the derivative of the stress tensor $\partial_j \Sigma_{ji}$ effectively contains the force density of the fluid, integration of this quantity over the volume of the particle yields the total force exerted on the particle. By taking into account the position $r_i$ at which each force element acts relative to the microparticle's center of mass, the torque density is $\in_{ilk}\, r_l \partial_j \Sigma_{jk}$ where $\in$ is the Levi-Civita symbol. Integration yields

$$F_i = \int_{\Omega} \partial_j \Sigma_{ji} d\Omega \tag{5}$$

$$T_i = \int_{\Omega} \in_{ilk}\, r_l \partial_j \Sigma_{jk} d\Omega \tag{6}$$

which can be simplified further by replacing the volume integral over $\Omega$ with a surface integral $\partial\Omega$ using Stokes' theorem,

$$F_i = \int_{\partial\Omega} n_j \Sigma_{ji} dS \tag{7}$$

$$T_i = \int_{\partial\Omega} \epsilon_{ilk} \, r_l n_j \Sigma_{jk} dS \tag{8}$$

For clarity, the surface element $d(\partial\Omega)_i$ is decomposed into a scalar area element $dS$ and a normal $n_i$ such that $d(\partial\Omega)_i = n_i dS$.

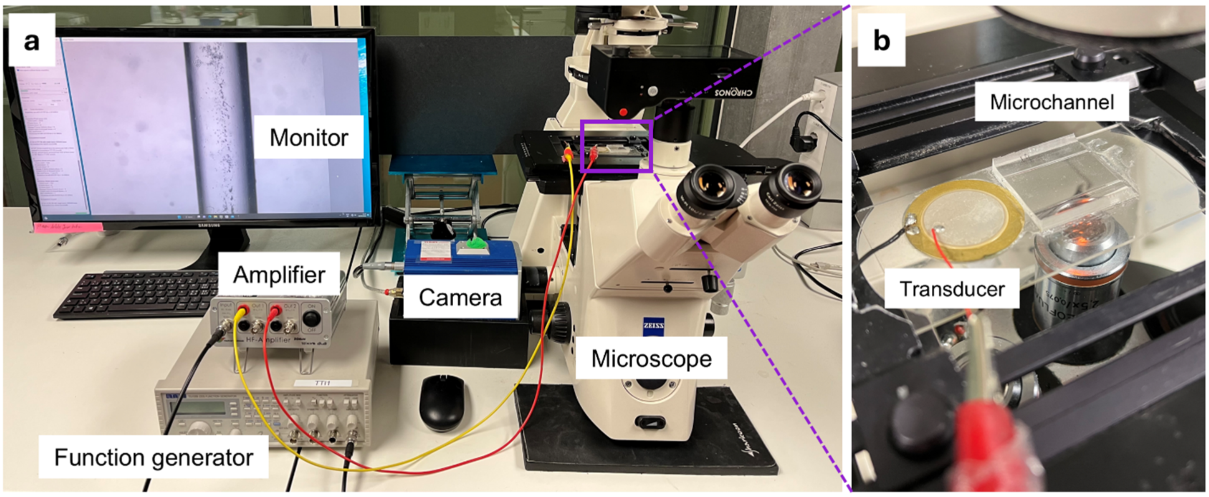


**Supplementary Fig. 1. Experimental setup for acoustic excitation and observation. (a)** Overview of the observation platform, illustrating the key components of the setup. **(b)** Detailed representation of the acoustic excitation device.

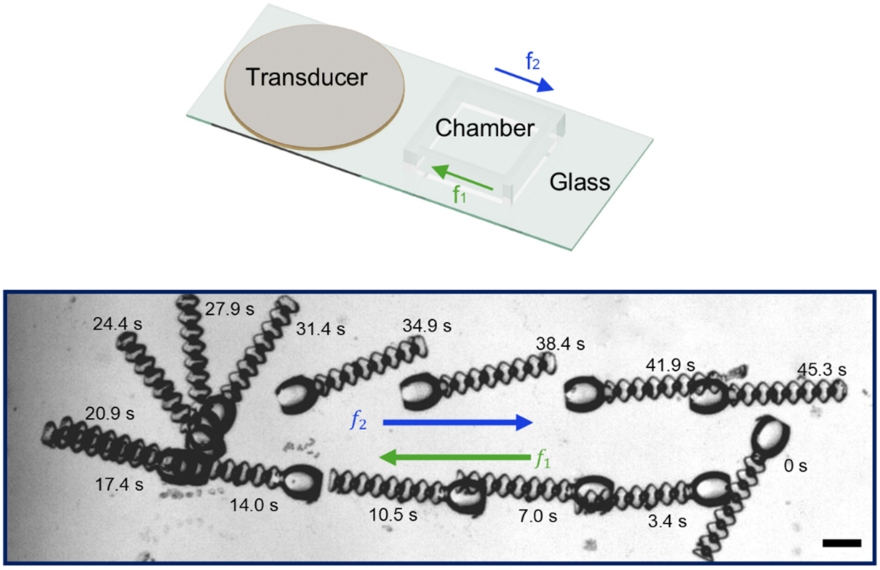


**Supplementary Fig. 2. Reorientation behavior of the microparticles in an open channel.** The head-helix microrobot also demonstrates bidirectional reorientation behavior in an open rectangular chamber, $f_1 = 8.8$ kHz, $f_2 = 13.4$ kHz, and 60 $V_{PP}$. Scale bars, 100 μm.

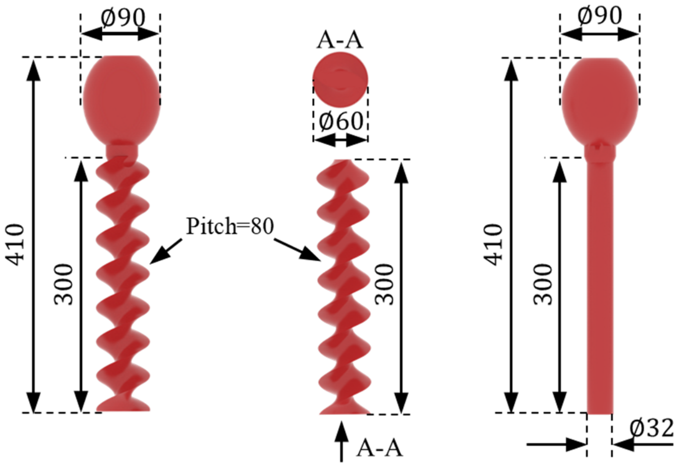


**Supplementary Fig. 3. Dimensions of the designed microparticles.** All dimensions in the schematic are given in micrometers.

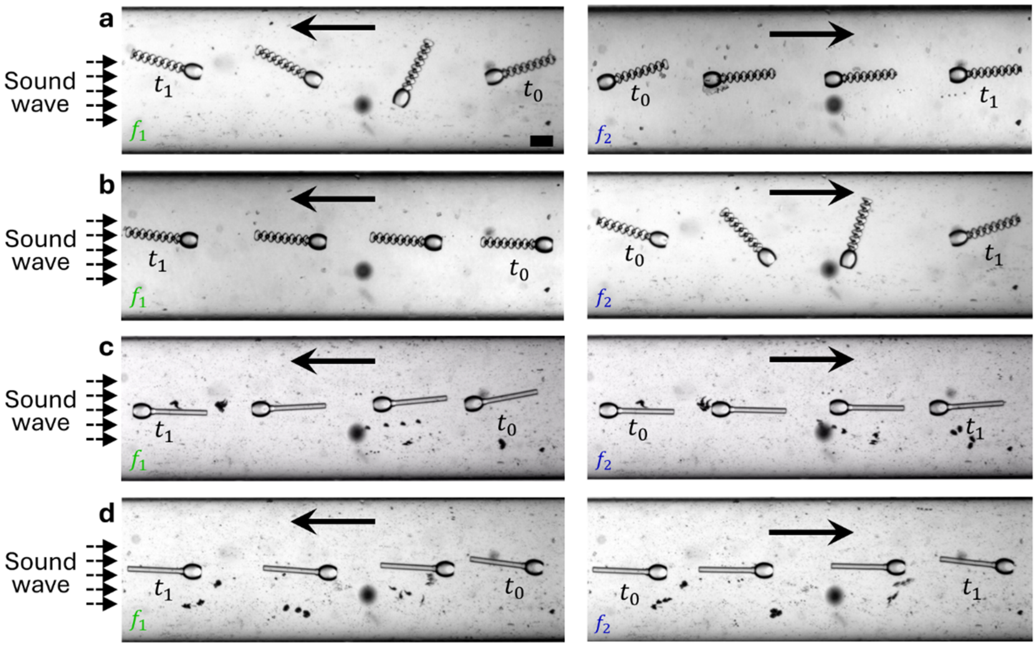


**Supplementary Fig. 4. The effect of initial orientations on microparticle movement.** (**a-b**) The movement behavior of a head-helix microparticle at $f_1$=12.3 kHz, $f_2$=8.8 kHz, and 60 $V_{PP}$: (**a**) $\theta = 0$ and (**b**) $\theta = \pi$. (**c-d**) The movement behavior of a head-cylinder microparticle at $f_1$=12.3 kHz, $f_2$=8.8 kHz, and 60 $V_{PP}$: (**c**) $\theta = 0$ and (**d**) $\theta = \pi$. Scale bar, 100 μm.

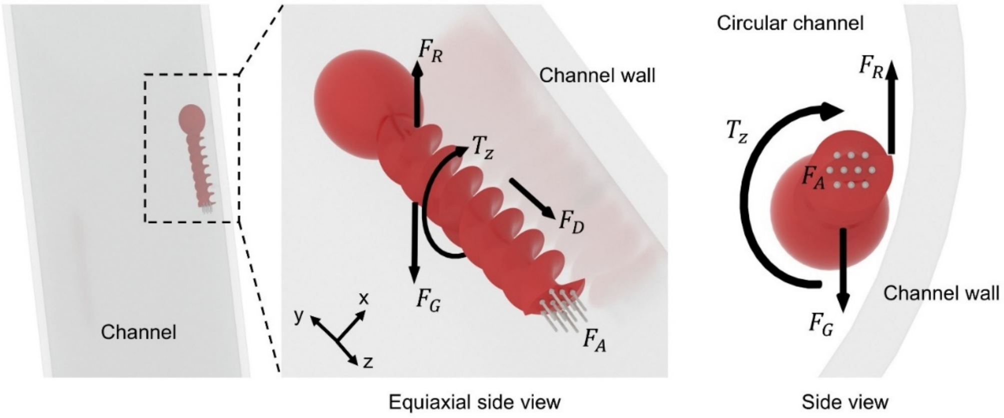


**Supplementary Fig. 5. Hydrodynamic model for an upstream-moving head-helix microparticle in a circular channel.** The particle is driven by acoustic radiation forces and a torque from tail-induced streaming, while wall interactions generate an additional torque and lateral forces. Balancing gravity $F_G$, acoustic radiation forces $F_A$, streaming-induced torque $T_Z$, resistive forces $F_R$, and drag forces $F_D$, the particle achieves unidirectional propulsion accompanied by continuous axial rotation along the channel wall.

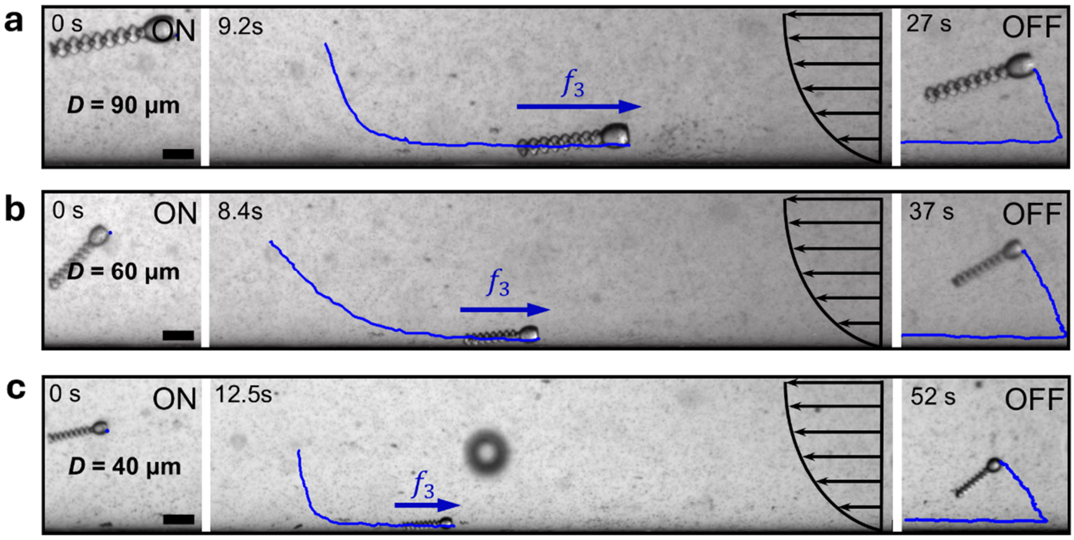


**Supplementary Fig. 6. Upstream motion of three particles with equal shape but different sizes (head diameters of *D* = 40 µm, 60 µm, and 90 µm).** Larger microparticles demonstrate faster velocities due to increased forces.

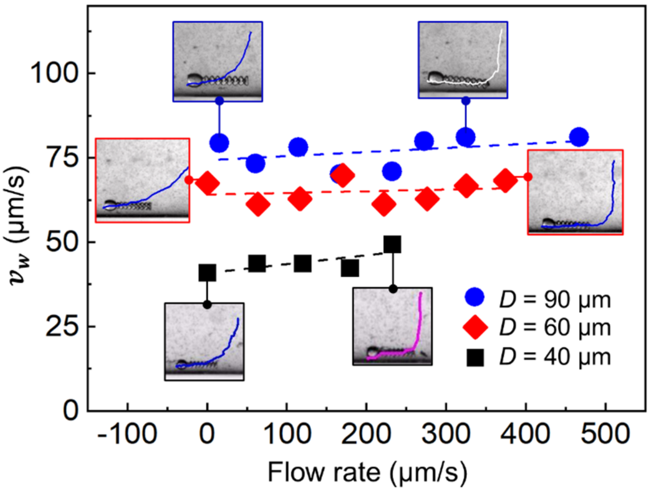


**Supplementary Fig. 7. Shift velocity of the microparticles.** The shift velocity represents the velocity of the three particles (*D* = 40 µm, 60 µm, and 90 µm) as they travel from the center bottom of the channel to the wall.
